# Natural motion statistics unify direction computations

**DOI:** 10.64898/2026.07.30.741322

**Authors:** Zixuan Deng, Wei Wei

**Affiliations:** Committee on Neurobiology, University of Chicago, Chicago, IL 60637, USA; Department of Neurobiology, Neuroscience Institute, University of Chicago, Chicago, IL 60637, USA

## Abstract

Visual direction selectivity is typically studied using coherent, unidirectional stimuli. However, natural vision involves more complex, heterogeneous motion fields generated by self-motion and moving objects. How visual circuits extract direction information from these motion fields is not fully understood. Here, we show that On-Off direction-selective ganglion cells (DSGCs) in the mouse retina encode an integrated motion direction from component motion vectors within their receptive fields. Using plaid stimuli with distinct velocity components, we demonstrate that DSGCs maintain unidirectional tuning even when component velocity vectors diverge by over 90 degrees. DSGCs’ preferred directions follow the angular mean, rather than the vector sum of motion vectors, a property emerging from the directional tuning of their inhibitory inputs. Intriguingly, while angular mean and vector sum represent distinct mathematical operations, the statistical properties of natural movies cause them to converge onto a single aligned direction, eliminating the necessity of the visual system to choose one operation over the other. We found that DSGCs encode this aligned direction for both optic flow and object motion movies, and the precision of direction encoding depends on the homogeneity of motion vectors in the receptive field. Our findings reveal that the retina performs an initial integration of motion vectors using angular averaging. Since natural motion statistics cause possible motion vector computations to converge, angular averaging effectively informs the animal about the consensus motion direction in natural environments.

## Introduction

The history of visual neuroscience is, in many ways, the history of parsing the complex visual world into experimentally tractable features^1,2^. If a feature could be defined — an orientation, a direction, a color — it could be isolated in a stimulus, systematically varied, and used to ask whether neurons are tuned to it. The answer, again and again, has been yes. Beginning with the discovery of orientation-selective neurons in cat visual cortex^1^, decades of work have revealed feature-selective neurons throughout the visual system across animal species^3,4^, establishing feature selectivity as the dominant framework for defining what visual circuits compute.

The vertebrate retina has been central in revealing not only what features the visual system computes^5^, but also the circuit mechanisms underlying feature selectivity^6,7^. Composed of three cellular layers and two synaptic layers, the retina transforms light patterns into parallel feature channels and relays feature information to downstream brain regions through the spiking outputs of retinal ganglion cells (RGCs). Each RGC type reports the output of a distinct neural circuit that utilizes multiple interneuron types with specific patterns of synaptic connections^8^. Together, these circuits enable RGCs to inform the rest of the brain about ethologically relevant visual information necessary for perception and behavior^9^.

Retinal direction selectivity provides one of the most elegant examples of how neural circuits give rise to feature selectivity. When presented with a moving stimulus, direction-selective ganglion cells (DSGCs) fire strongly to motion along their preferred directions and minimally to motion in the opposing, null directions^10^. This computation critically depends on direction-selective inhibition from starburst amacrine cells (SACs)^11–13^, inhibitory interneurons whose radially symmetric dendrites are themselves tuned to motion outward from the soma^14^. This outward direction selectivity of the SAC is then conveyed to DSGCs by selective wiring: each SAC dendritic quadrant preferentially inhibits a specific DSGC subtype, giving rise to the DSGC’s direction-selective spiking responses^12,15–18^. This canonical SAC-DSGC circuit motif is responsible for the direction selectivity of both On and On-Off classes of DSGCs^16,18^. On-DSGCs respond selectively to light increments, are tuned to slow global motion, and project to the accessory optic system (AOS) to drive image-stabilizing reflexes^9,19–22^; On-Off DSGCs respond to both light increments and decrements, maintain robust tuning across a broad range of velocities, and project to image-forming brain regions including the superior colliculus (SC) and dorsal lateral geniculate nucleus (dLGN) of the thalamus^23–28^.

Despite this detailed circuit-level understanding, it remains unclear how DSGCs extract motion direction from natural scenes. Unlike simple artificial motion stimuli, natural scenes usually contain dynamic and heterogeneous velocity components^29^. Whether motion comes from the gaze of a cat tracking prey, or from a mouse fleeing its predator, object and self-motion can occur simultaneously and move in different directions, producing heterogeneous motion vectors within a single cell’s receptive field that vary over time. This raises a fundamental question for the direction-selective circuit: when multiple motion vectors are present in the receptive field, which direction does a DSGC encode?

In this study, we address the above question using both artificial and natural motion stimuli. We found that for both types of motion stimuli, the preferred direction of the On-Off DSGC largely follows the angular mean direction of local motion vectors. The amplitude of the direction signal increases with temporal contrast, while the precision of direction encoding decreases as motion vectors become more dispersed. We further show that these changes in DSGCs’ tuning properties arise from inhibitory inputs from SACs. Remarkably, despite the complexity of natural motion, distinct vector operations of vector sum and angular mean converge onto a unified direction, thereby simplifying the problem of direction extraction from natural scenes. Together, our results indicate that natural motion statistics provide a shortcut for retinal direction computation: angular averaging is sufficient to extract a unified motion direction that can inform the animal about movement in the natural environment.

## Results

### On-Off DSGCs integrate component vectors in Type II plaids in a cross-angle dependent manner

To investigate how On-Off DSGCs integrate motion vectors within their receptive fields, we first used a two-vector artificial motion stimulus, the Type II drifting plaid^30^, which consists of two superimposed drifting gratings with identical spatial frequency and contrast but different speeds (11°/s and 33°/s) and directions (cross angles between the two component gratings: 15°, 45°, 90°, 135°, 160°) (Fig. 1a and 1b). On-Off DSGC responses were measured by two-photon calcium imaging in a transgenic mouse line carrying Vglut2-Cre+ and floxed GCaMP6f, which specifically labels retinal ganglion cells in the mouse retina^31^ (Extended Data Fig. 1a and 1b). On-Off DSGCs were identified by their direction-selective responses to individual component drifting gratings (Fig. 1c, top two rows and Methods) and to both the leading and trailing edges of a moving bar stimulus (Extended Data Fig. 1c and Methods). We quantified tuning strength using the global direction selectivity index (gDSI), which is the magnitude of the normalized complex sum of responses to stimuli moving in different directions (Methods). On-Off DSGCs show similar directional tuning to both the slow (11°/s) and fast (33°/s) gratings (Extended Data Fig. 1d), though slower gratings evoked a slightly greater gDSI and peak amplitude (Fig. 1c, top two rows, and Extended Data Fig. 1d and 1e). In response to Type II plaids, On-Off DSGCs exhibit unimodal directional tuning curves even at obtuse cross angles, indicating that they signal one integrated motion direction from the two component motion vectors (Fig. 1c and Extended Data Fig. 1f).

**Fig. 1.**
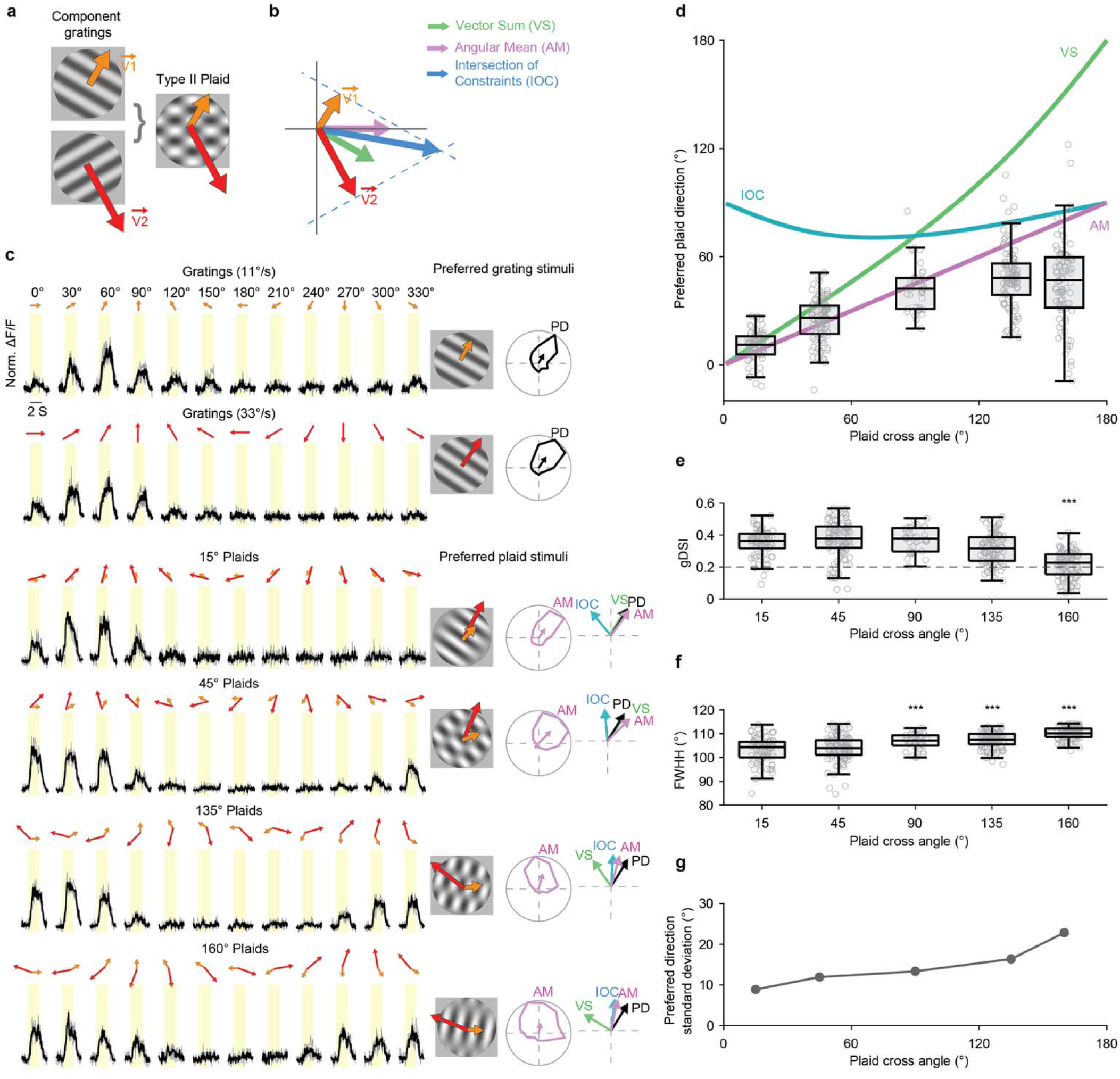
On-Off DSGCs’ directional tuning varies with the cross angle of component vectors in Type II plaids. **a**, Schematic of Type II plaid composed of two drifting gratings separated by 120° cross angle and 1:3 speed ratio. **b,** Three distinct vector computations for plaid motion direction: VS, AM, and IOC. IOC is determined by the intersection of the dashed lines perpendicular to V1 and V2. **c,** Left, calcium responses of an example DSGC to gratings (11°/s and 33°/s) and plaids. Columns indicate stimulus direction; yellow shading marks stimulus duration. Gray traces show single trials and black traces show trial averages; *n* = 3 trials. Middle, preferred grating and plaid stimuli for the same cell. Right, grating tuning curves plotted in black with normalized vector sum. Plaid tuning curves in magenta are plotted against the AM direction. To the right of the plaid tuning curves are reference directions: a cell’s preferred direction, PD, black arrow, is determined by drifting gratings. If a cell prefers plaid AM, the magenta AM arrow should align with PD. **d,** Population plaid PD as a function of plaid cross angle. **e,** Global direction selectivity index (gDSI) at different plaid cross angles (One-way ANOVA, *P* < 0.01; Tukey-Kramer post hoc comparisons were made against 15° plaids: 45° versus 15°, *P* = 0.74; 90° versus 15°, *P* = 0.86; 135° versus 15°, *P* = 0.10; 160° versus 15°, *P* < 0.001). **f,** Tuning width at different plaid cross angles. (One-way ANOVA, *P* < 0.001; Tukey-Kramer post hoc comparisons were made against 15° plaids: 45° versus 15°, *P* = 0.75; 90° versus 15°, *P* < 0.001; 135° versus 15°, *P* < 0.001; 160° versus 15°, *P* < 0.001). **g,** Circular standard deviation of plaid PDs as a function of plaid cross angle. In **d–f**, open circles indicate individual cells; center lines indicate medians; boxes indicate interquartile ranges (IQRs); whiskers indicate 1.5 × IQR for this and subsequent figures. In **d–g**, *n* = 76, 99, 37, 94 and 93 cells for 15°, 45°, 90°, 135° and 160° plaids, respectively.

To determine how On-Off DSGCs integrate component vectors, we compared each cell’s preferred direction to drifting gratings with that to Type II plaids (Fig. 1c). The motion direction of Type II plaids can be quantified using three candidate integration rules: vector sum (VS), angular mean (AM), and intersection of constraints (IOC)^32^ (Fig. 1b). VS is the mathematical vector addition of the two velocity vectors, which biases the resultant direction toward the faster grating. AM is the mean angle of the two component directions regardless of speed. IOC is defined by the intersection of the two component constraint lines perpendicular to each velocity vector, which corresponds to the motion direction of the plaid pattern^30^. We found that the preferred directions of On-Off DSGCs to Type II plaids depend on the cross angle of the component gratings (Fig. 1c and 1d). At acute cross angles (15° and 45°), On-Off DSGCs are strongly tuned to the AM direction. As the cross angles increase to the obtuse range, the same cells’ preferred directions deviate from the AM direction toward the direction of the slower grating (Fig. 1d) and become more variable (Fig. 1d and 1g). At 160° cross angle, we observed a significant decrease of gDSI compared to the 15° cross angle (Fig. 1e), and an increased tuning width (Fig. 1f).

These results indicate that at acute cross angles of Type II plaids, On-Off DSGCs are reliably tuned to the AM direction. However, as cross angle increases, both the tuning strength and the precision of their direction selectivity decrease.

### Cross-angle dependent tuning of inhibitory inputs dictates On-Off DSGCs’ directional tuning to Type II plaids

Since the direction selectivity of On-Off DSGCs primarily arises from directionally tuned inhibitory inputs from SACs^11–13^ (Fig. 2a and 2b), we next investigated whether DSGC inhibitory postsynaptic currents (IPSCs) exhibit the same cross-angle-dependent directional tuning as seen in their spiking outputs. We performed two-photon targeted voltage clamp recordings from On-Off DSGCs preferring the posterior direction using Drd4-eGFP mice^23^ (Methods) to isolate their synaptic inputs during 15°, 90°, and 160° plaids. For each cell, we first determined the IPSC preferred direction using slow and fast gratings (Fig. 2b and 2c). We observed that slow and fast gratings evoked similar IPSC peak amplitude and tuning (Fig. 2b and 2c), consistent with previous work showing that GABAergic inhibition is largely speed invariant^21,33^. We then determined the IPSC preferred directions for Type II plaids of different cross angles, and compared these measured preferred directions with the VS, AM, and IOC directions of the preferred plaid stimuli (Fig. 2d). IPSC tuning closely mirrored the cross-angle dependent tuning of the ganglion cell calcium responses (Fig. 2e): at smaller cross angles (15° and 90°), inhibitory preferred directions clustered near the AM direction, but at an obtuse cross angle (160°) they deviated from the AM direction. At 160° plaids, we also observed reduced direction selectivity (Fig. 2f) and increased variability in preferred direction angles (Fig. 2g). Thus, these results indicate that inhibitory input is sufficient to account for the cross angle-dependent directional tuning properties of On-Off DSGCs.

**Fig. 2.**
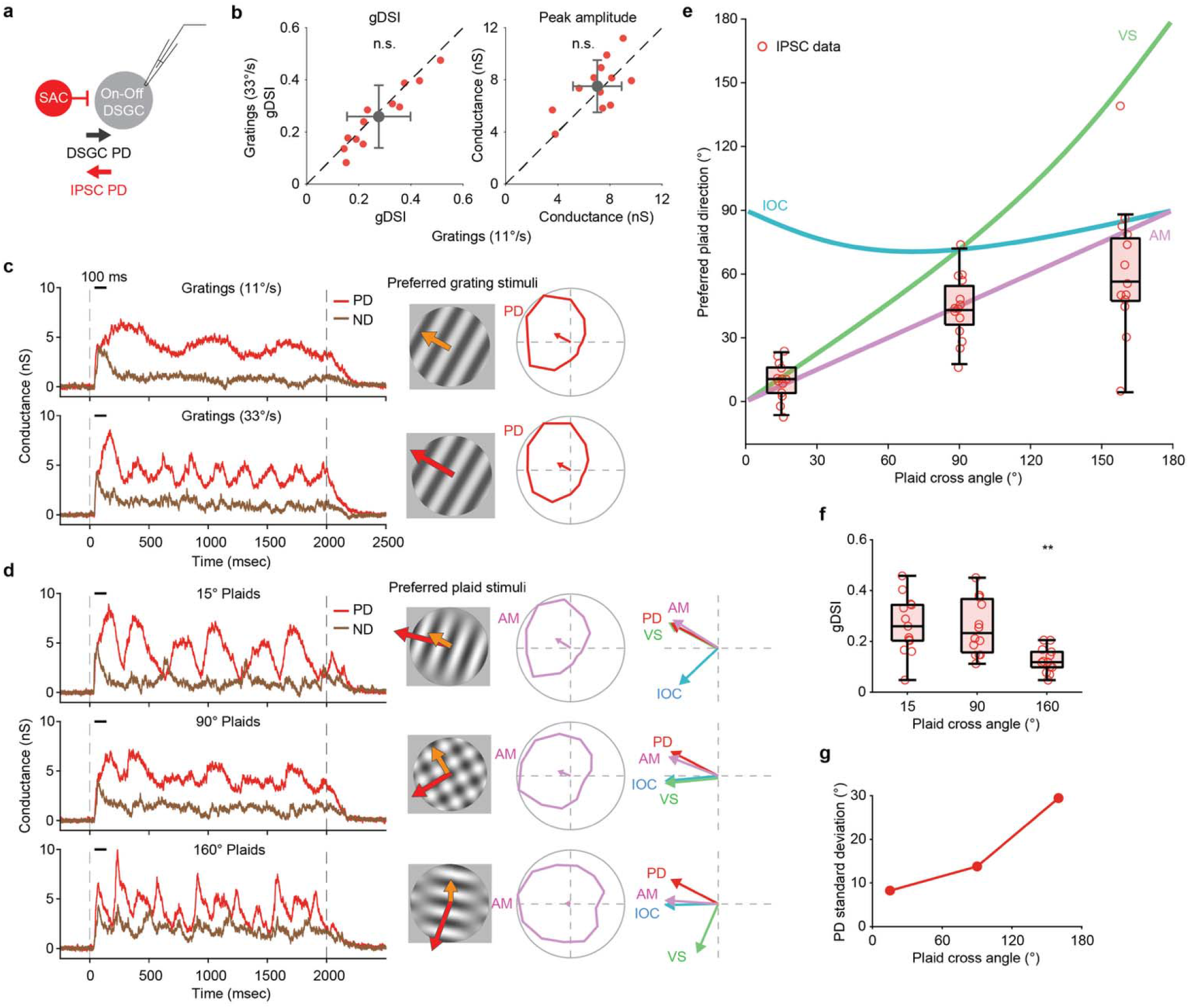
DSGC inhibitory tuning varies with plaid cross angle. **a**, Simplified schematic showing that starburst amacrine cells (SACs) provide directionally tuned inhibition to an On-Off DSGC. **b**, Comparison of IPSC tuning evoked by slow (11°/s) versus fast (33°/s) drifting gratings. Red points show individual cells; dashed lines indicate unity; gray symbols and error bars indicate mean ± s.d. IPSC gDSI and peak amplitude do not differ significantly between grating speeds (*n* = 12 cells; gDSI, *P* = 0.15; peak amplitude, *P* = 0.30, Wilcoxon signed-rank test). **c**, Left, inhibitory conductance of an example On-Off DSGC to slow (11°/s) and fast (33°/s) gratings. Traces show responses for the preferred direction (PD, red) and null direction (ND, brown). Dashed vertical lines indicate stimulus onset and offset. Middle, preferred gratings stimuli. Right, corresponding IPSC tuning for the same cell. **d**, Left, inhibitory conductance from the same cell in **c** to Type II plaids with cross angles of 15°, 90° and 160°. Middle, preferred plaid stimuli for the same cell. Right, plaid tuning curves in magenta are plotted against the AM directions. If inhibition prefers the AM direction, the magenta arrow should align with the red PD arrow. **e**, Population IPSC preferred plaid direction as a function of plaid cross angle. *n* = 13, 14 and 13 cells for 15°, 90° and 160° plaids, respectively. Directions of different mathematical vector computations are indicated by curves of different colors. **f**, Tuning strength of DSGC inhibition varies with plaid cross angle. One-way ANOVA, *P* < 0.001. Tukey–Kramer post hoc tests showed lower gDSI at 160° than at 15° (*P* < 0.01) or 90° (*P* < 0.01), with no difference between 15° and 90° (*P* = 0.98). *n* values are the same as in **e**. **g,** Circular standard deviation of IPSC preferred direction as a function of plaid cross angle. *n* values are the same as in **e** and **f**.

### Temporal contrast sets the time course of DSGC excitatory inputs during Type II plaids

We next investigated the tuning of excitatory postsynaptic currents (EPSCs). EPSCs were weakly tuned to motion direction for plaids of all cross angles (Fig. 3b and 3c). However, we found distinct and stereotyped EPSC waveforms for each cross angle, indicating that the plaid texture activated excitatory inputs in distinct spatiotemporal patterns (Fig. 3b). Because DSGCs pool glutamatergic inputs from an array of On and Off bipolar cells with smaller receptive fields^34–37^ (Fig. 3a), we next asked whether the temporal contrast of the stimulus at individual bipolar cell RF underlies the distinct temporal structure of DSGC excitation. We computed temporal contrast as the frame-to-frame luminance change, half-wave rectified into On and Off components, and averaged across subunits (Methods). Interestingly, local pooling of rectified luminance signals across On and Off bipolar cell RFs (2° diameter bipolar RF units with 1° offsets across a 7° diameter DSGC RF) better recapitulated the time course of DSGC excitatory inputs than luminance averaged across the entire DSGC receptive field for all plaid cross angles (Pearson *r* for bipolar cell subunit pooling versus receptive-field averaging: 0.74 versus 0.72 at 15°, 0.91 versus 0.85 at 90°, and 0.78 versus 0.69 at 160°; Fig. 3b and 3d; Extended Data Fig. 2a; Methods). These results indicate that bipolar cell-level temporal contrast pooling sets the time course of DSGC excitation during Type II plaids.

**Fig. 3.**
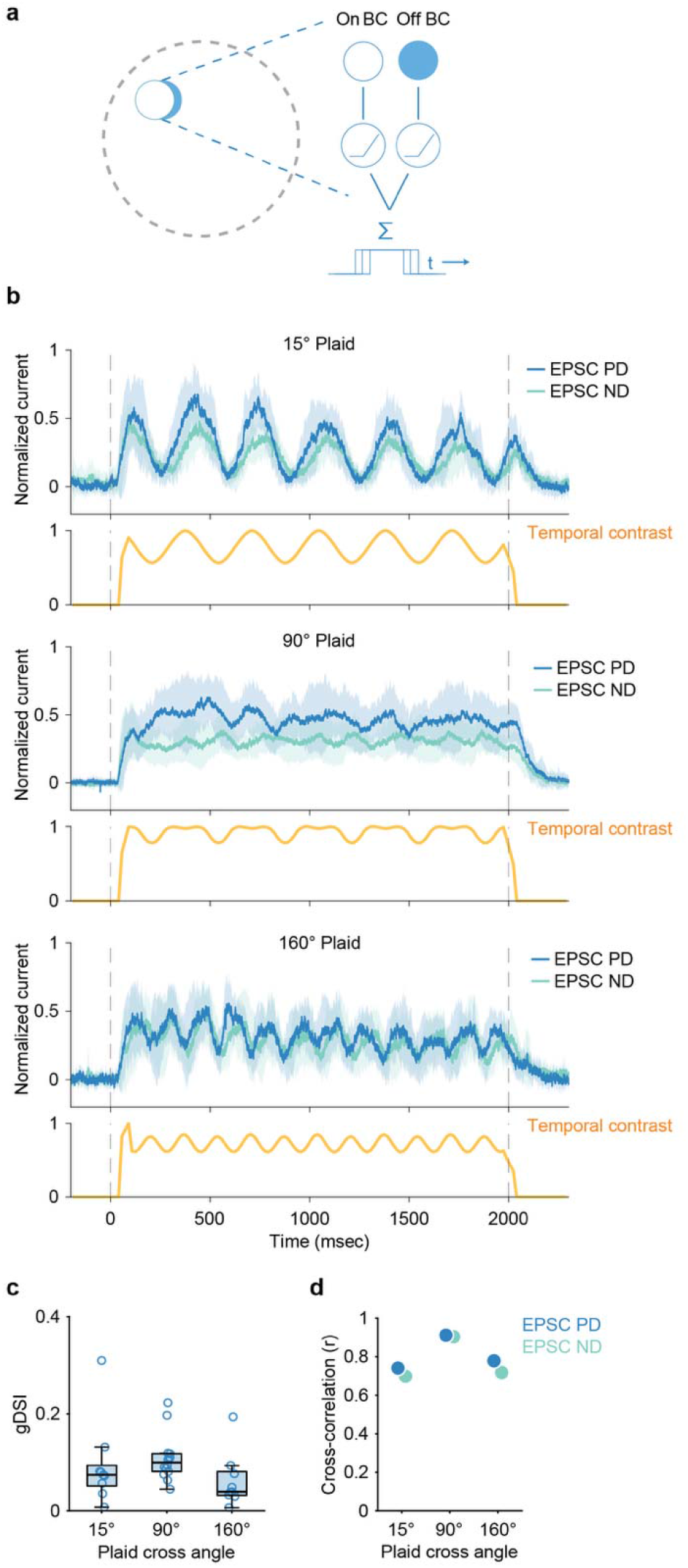
Excitatory inputs onto DSGCs track stimulus temporal contrast for Type II plaids. **a**, Schematic illustrating DSGC pooling of local ON and OFF bipolar-cell inputs within its receptive field. **b**, Normalized EPSCs evoked by drifting plaids at cross-angles of 15°, 90° and 160°. For each cell, the preferred direction (PD) and null direction (ND) were assigned from the plaid-evoked IPSC tuning measured at the same cross-angle. Traces show mean ± s.d.; *n* = 9, 13 and 9 cells, respectively. Dashed lines mark stimulus onset and offset. Normalized temporal contrast, shifted by 40 ms to account for response latency, is shown below each trace. **c**, EPSC gDSI across plaid cross-angles. EPSC direction selectivity was weak and did not differ across cross-angles; one-way ANOVA, *P* = 0.25. *n* = 9, 14 and 9 cells for 15°, 90° and 160° plaids, respectively. One cell in the 90° condition did not have IPSC tuning and was therefore excluded from the response traces in **b**. **d**, Cross-correlation between stimulus temporal contrast and mean PD or ND EPSC responses.

Together, synaptic currents measured during Type II plaids support a circuit model in which excitation provides a temporal gate on DSGC activation through local pooling of stimulus temporal contrast, and inhibition sets DSGC directional tuning by integrating the component vectors in a cross-angle-dependent manner.

### Quantification of motion vectors in natural movies

Having established that the direction-selective circuit integrates the two motion vectors in the plaid stimulus in a cross-angle-dependent manner, we next investigated the implications of this finding for DSGC motion signaling in natural scenes. The first step is to analyze motion vectors in natural movies. We focused on two types of natural motion stimuli: optic flow driven by self-movement, and object motion from external movement (Fig. 4a; Methods). For optic flow, we analyzed outdoor footage from an open-source database taken from the mouse’s perspective (a few centimeters from the ground) using a fisheye lens^38^. We selected four segments (20 s each) from different outdoor environments in the database. For object motion, we analyzed phone recorded videos featuring cats^39^, mice, squirrels and birds to include diverse movement motifs: swatting, turning, grooming, jumping, and flying (Extended Data Fig. 3). In these recordings, the camera was largely stationary, with occasional panning consistent with natural shifts in viewpoint. Unlike optic flow, which produces dense and continuous motion, object motion depended on the animal’s behavior in the movie: it was largely confined to the animal in motion and punctuated by frequent pauses. We sampled 16 short clips (∼7-10 s each) to include movements that are behaviorally salient to mice, including those of conspecifics and potential predators. Although the selected optic flow and object motion movies represent distinct behaviors of the observer, active navigation versus stationary observation, the motion fields in each movie contain elements of both optic flow and local motion, with one dominating. For example, in optic flow movies, wind blows on the grass causing local movement; in cat movies, optic flow occurs during camera panning. We did not include movies featuring only global image shift during rapid saccadic eye movements ^40,41^.

**Fig. 4.**
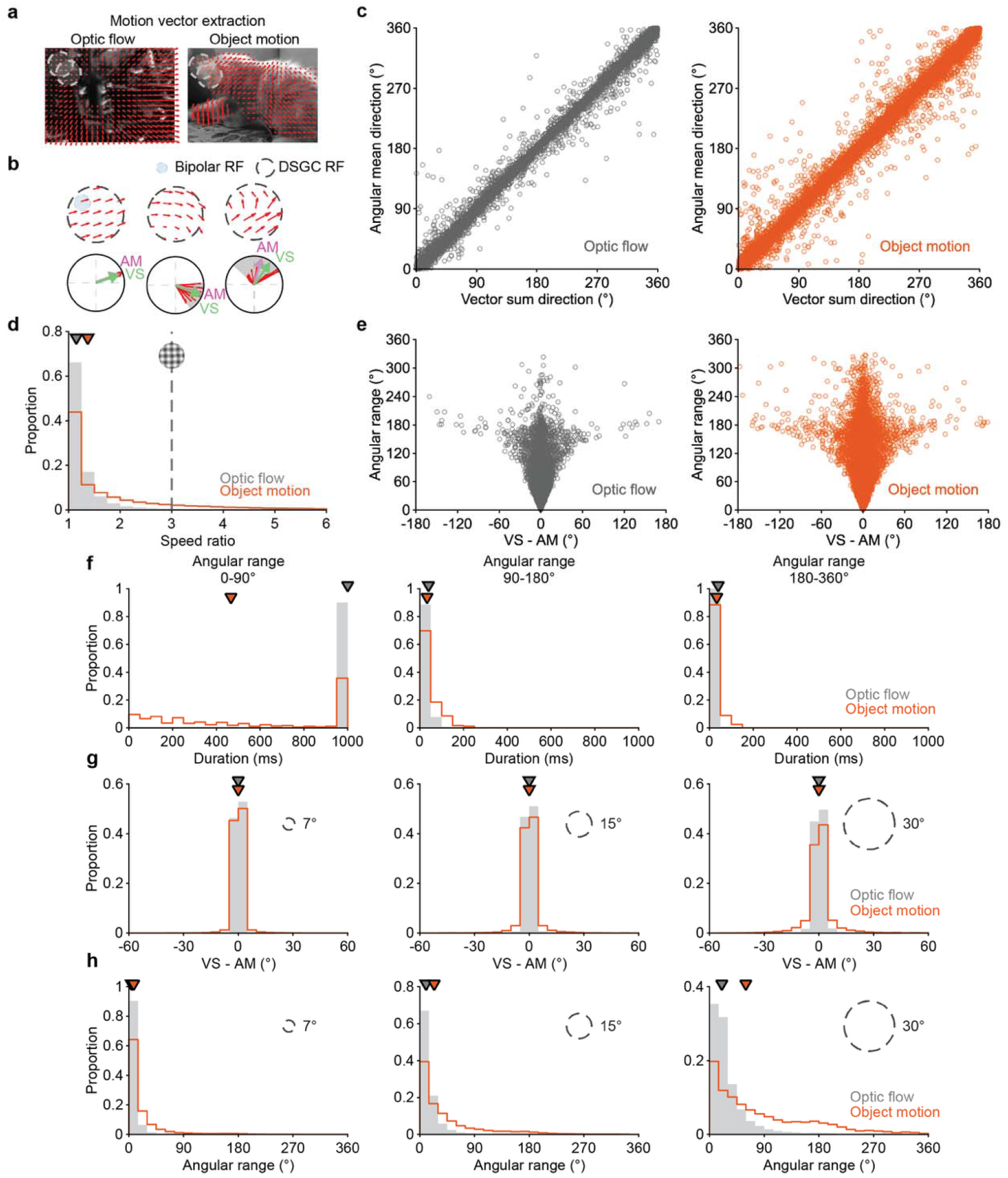
VS and AM directions of motion vectors during optic flow and object motion converge to a consensus direction. **a**, Example motion fields extracted from natural movie frames using RAFT. Red arrows indicate local motion vectors; the gray dashed circle indicates the DSGC receptive field. **b,** Top, three examples of motion vector distributions within a DSGC RF; each vector corresponds to motion within a bipolar-cell RF. Bottom, corresponding angular range (gray shaded wedges), VS directions (green), AM directions (magenta), and component motion vectors (red) for the above examples with angular ranges of 8°, 63°, and 109°. **c**, Comparison of VS and AM directions across all DSGC RFs and movie frames for optic flow (left) and object motion (right). Each point represents one RF across consecutive frames. VS and AM were highly correlated for optic flow and object motion (*r* = 0.99 for both). Object-motion statistics were computed from 16 movie clips, 3,894 frames and 100,077 sampled RFs; optic-flow statistics were computed from 4 movie clips, 2,002 frames and 312,518 sampled RFs. **d**, Distribution of speed ratios within sampled RFs. Median ratios were 1.15 for optic flow and 1.37 for object motion. The dashed vertical line indicates the speed ratio of 3 for Type II plaids. **e**, Scatterplot between VS- AM offset and angular range across sampled frames. Larger angular ranges are associated with greater variability in VS-AM offset. **f**, Distributions of movie durations when local vectors moved within different angular range bins. For 0-90°, median durations were 1000 ms for optic flow and 466 ms for object motion. For both 90-180° and 180-360°, median durations equaled one frame interval (∼40 ms) for optic flow and object motion. **g,** Distributions of VS–AM offsets computed from the same optic-flow and object-motion movie datasets analyzed in **c-f**, shown for RF diameters of 7°, 15° and 30°. Median VS– AM offset remained close to 0 across RF diameters. **h**, Distributions of angular ranges for the same RF diameters and movies as in **g**. Median angular ranges were 4°, 10°, and 20° for optic flow, and 8°, 24°, and 60° for object motion at 7°, 15°, and 30° RF sizes, respectively. For **c–h**, gray denotes optic flow and orange denotes object motion. Inverted triangles in **d, f, g** and **h** indicate medians.

To quantify the component motion vectors within individual DSGC receptive fields, we first computed pixel-wise motion fields in natural movies using RAFT^42^, a state-of-the-art deep recurrent neural network that directly correlates pairs of pixels across consecutive frames (Methods), which improves sensitivity to small, fast-moving objects that can be missed by coarse-to-fine pipelines such as the Farneback method^43^. We then parsed vectors in these motion fields into a bipolar cell RF mosaic spanning a DSGC receptive-field center (2° diameter bipolar RF units with 1° offsets across a 7° diameter DSGC RF) by averaging pixel-wise motion vectors within each bipolar cell’s RF (Fig. 4b and Methods). Motion vectors with 5°/s - 60°/s speed range, in which On-Off DSGCs show robust responses^21,44,45^, are used in the calculation. Together, this method yields a circuit-constrained estimate of the component motion vector ensemble that is most likely to modulate DSGC firing.

### Vector sum and angular mean directions converge for natural stimuli

We next calculated the vector sum (VS) and the angular mean (AM) directions within the DSGC RF across natural movie frames (Fig. 4b). Unlike with Type II plaids, the IOC direction is not always well-defined because natural scenes do not solely consist of a single rigidly moving pattern. Therefore, the constraint lines of component vectors do not converge on a single intersection point. More importantly, the VS and AM directions were closely aligned for natural movies. For Type II plaids, VS and AM are distinct functions of vector cross angles (see green and magenta lines in Fig. 1d). In sharp contrast, for both optic flow and object motion movies, the VS and AM converge to the same direction (Fig. 4c). The alignment was present in every clip analyzed, across all movement motifs sampled (per-clip Pearson *r* = 0.97-1.00; Extended Data Fig. 3), demonstrating that these two vector computations describe the same motion direction in natural scenes.

Next, we asked which features of natural motion statistics could account for the striking alignment between VS and AM directions. Because motion vectors were restricted to speeds of 5°/s - 60°/s in the above analysis, we then tested whether this speed filtering step imposed the observed alignment. We first examined the distribution of pixel-wise motion speeds. As expected, natural movies contained substantial static regions and were dominated by slow speeds^46,47^ (Extended Data Fig. 4a). We then tested the extreme case of applying no speed filter and calculated VS and AM directly from motion vectors in pixels per frame within DSGC receptive fields throughout each movie. VS-AM alignment persisted without converting pixel displacement to angular velocity or imposing any speed thresholds (Extended Data Fig. 4b and 4c).

We next sought to understand why the VS-AM alignment is so robust in natural movies. Because VS– AM divergence depends on the relative speed of component vectors, we quantified the speed ratio, defined as the ratio between the maximum and minimum speed of component vectors within the DSGC RFs. We found that although larger speed ratios are more likely during object motion than optic flow, the median speed ratios of both are near one: 1.15 for optic flow and 1.37 for object motion (Fig. 4d). Thus, larger relative speed differences between component vectors are uncommon in natural movies, providing one explanation for the close alignment of the VS and AM directions.

Because the VS–AM differences for Type II plaid stimuli were most pronounced with large angular separations (Fig. 1d), we next examined whether a similar relationship applies to natural movies. To quantify the dispersion of component vectors in natural movies, we used the angular range, which describes the minimal circular interval spanning all component vector directions (Fig. 4b). We then quantified the relationship between VS–AM difference and angular range (Fig. 4e). When angular range was narrow (<90°), VS and AM directions were closely aligned. As angular range increased, the difference between VS and AM directions became more variable (Fig. 4e). However, these broad angular range epochs (>90°) were rare and almost never persisted beyond a single movie frame in either optic flow or object motion (Fig. 4f). Thus, instances in which the VS and AM were likely to diverge typically occurred only briefly across adjacent movie frames and were unlikely to persist over the temporal integration window of DSGC direction selectivity (approximately 100 ms, based on the minimal time for preferred and null IPSCs to diverge, Figs. 2c and 2d). Overall, coherent motion fields dominate natural scenes, even during object motion. These results indicate that the convergence between VS and AM directions in natural movies reflects the spatiotemporal structure of the complex motion vector ensemble.

To further test whether the alignment of VS and AM directions in natural scenes depends on receptive field size, we computed the difference between VS and AM directions for the same set of movies while varying the size of the DSGC RF diameter (Fig. 4g). As we increased the RF size, angular range increased for both object motion and optic flow (Fig. 4h). Strikingly, we found that VS and AM directions remained aligned as RF diameter increased from 7° (DSGC RF size) to 15° (RF size of superficial SC and V1^48^) to 30° (RF size of visual areas downstream of V1^49^). Together, these results indicate that vector computations that are distinct for Type II plaids remain closely aligned in natural scenes across the RF diameters tested.

### Natural movie’s temporal contrast modulates DSGC preferred-direction firing rate

To probe On-Off DSGC responses to natural movies, we performed two-photon targeted spike recordings in Drd4-eGFP mice^23^ in response to one representative optic-flow movie and an object motion movie with distinct motion statistics (Fig. 5a and b; see also Supplementary Video 1 and 2). We cropped each natural movie to a 10° diameter aperture, slightly larger than the 7° diameter DSGC average receptive-field center, to ensure that the movie covered the receptive-field center of every recorded cell despite cell-to-cell variation in center size (Methods). The optic flow movie features grass moving with highly coherent motion trajectories whereas the object motion movie features a cat paw swatting a ball, which exhibited markedly different motion statistics (Fig. 5b, 5c and 5d). For each cell, we first measured spiking responses to drifting gratings in 12 directions to estimate the preferred direction, and then measured responses to the natural movies rotated in 12 orientations (0°–330° in 30° increments), with two repetitions per orientation (Fig. 5e and Methods).

**Fig. 5.**
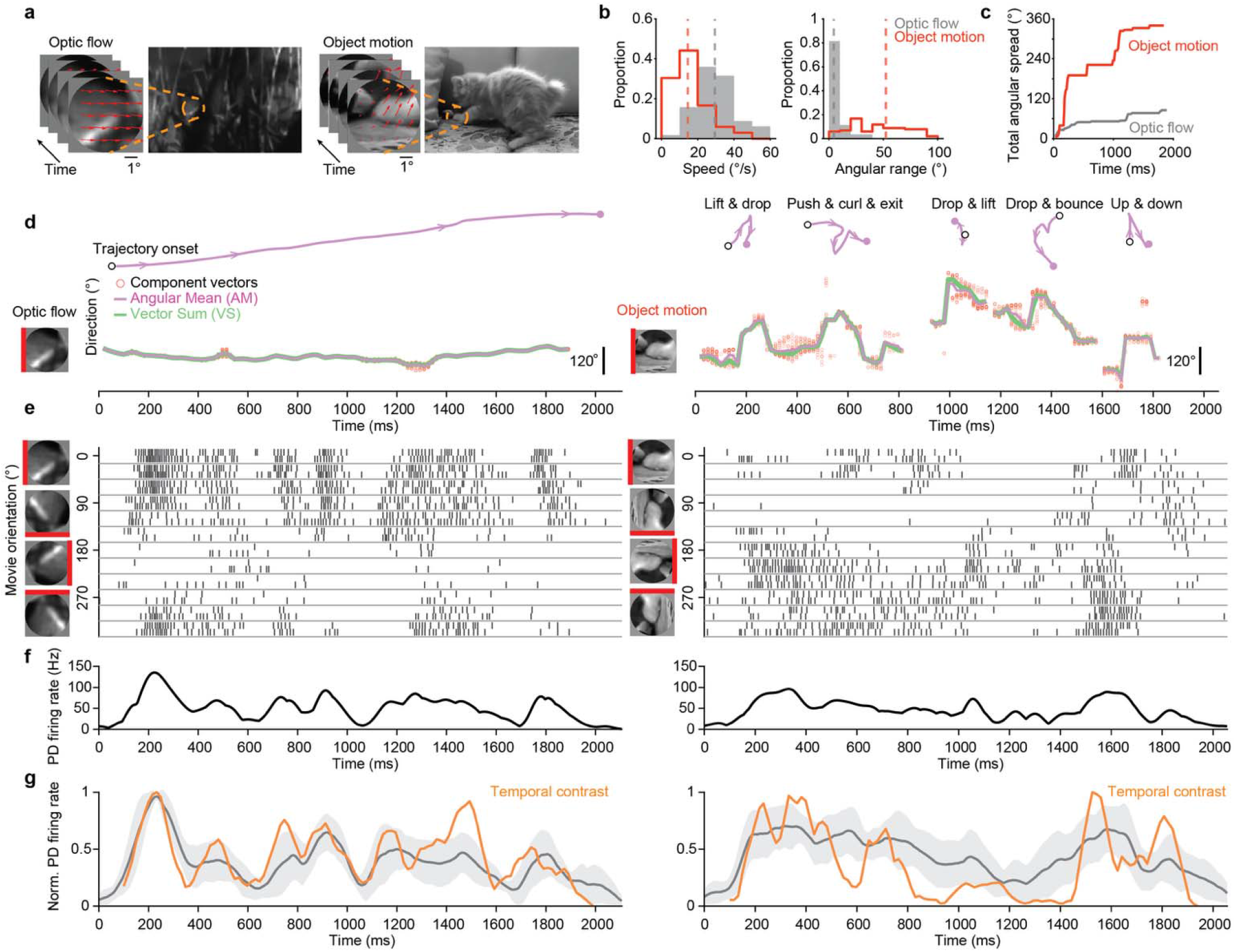
Natural movie temporal contrast modulates DSGC preferred direction firing. **a**, Example frames from optic- flow and object-motion movies used for visual stimulation. Orange dashed circles indicate the circular crop presented to the retina; red arrows indicate local component motion vectors within the cropped region. **b**, Motion statistics for cropped optic- flow and object-motion movies. Histograms show the distributions of frame-wise speeds and angular ranges. Median speed was 29°/s for optic flow and 15°/s for object motion; median angular range was 4° for optic flow and 52° for object motion. **c**, Cumulative angular spread across each movie. Total angular spread was 86° for optic flow and 340° for object motion. **d**, Motion direction of optic-flow and object-motion movies over time. Top, motion trajectory reconstructed from the frame- wise angular mean (AM) direction and mean speed; arrows indicate motion direction. Bottom, component motion vectors (red open circles) plotted over time, with the angular mean (AM, magenta) and vector sum (VS, green) overlaid. The object- motion movie comprises distinct movement bouts with intervening pauses during which no motion was detected. **e**, Example spike responses from the same DSGC to optic-flow (left) and object-motion (right) movies. Spikes are sorted by movie orientation, with two repetitions stacked for each orientation. **f**, Preferred-direction firing rate over time for the example cell in **e**, averaged across repetitions and smoothed with a 30-ms Gaussian kernel. **g**, Population preferred-direction firing rate and movie temporal contrast over time. Gray traces show mean normalized preferred-direction firing rate across DSGCs; gray shaded regions indicate standard deviation. Orange traces show normalized temporal contrast computed from the movie frames. *n* = 10 cells for optic flow and *n* = 15 cells for object motion. Firing rate was matched to each movie frame after applying a 100 ms latency shift. Temporal contrast was strongly correlated with preferred-direction firing rate for optic flow (Pearson *r* = 0.74, *P* < 0.001, *n* = 115 frames) and object motion (Pearson *r* = 0.67, *P* < 0.001, *n* = 112 frames).

Both optic flow and object motion evoked robust firing whose strength varied systematically with stimulus rotation (Fig. 5e). To capture the temporal structure of preferred-direction firing over time, we applied a sliding time window to calculate the maximum firing rate for each cell across the 12 stimulus rotations (Fig. 5f; Methods). We found that the dynamics of the preferred-direction response are well correlated with the temporal contrast of the stimulus, calculated either from luminance averaged from the entire RF (optic flow: Pearson *r* = 0.74; object motion: 0.67, Fig. 5g) or from local pooling of rectified luminance signals across On and Off bipolar cell (optic flow: Pearson *r* = 0.37; object motion: 0.56; Extended Data Fig. 2b). Together, our results indicate that the temporal contrast of natural movies modulates the amplitude of DSGC preferred direction firing.

### DSGCs are tuned to the aligned VS/AM direction during natural movies

We next asked which direction DSGCs prefer during optic flow and object motion. Unlike plaids, in which the component motion vectors remain fixed, motion vectors in natural movies change over time. We therefore used a sliding-window analysis (100 ms window, 20 ms step; Methods) to identify direction-selective windows for each cell across the stimulus duration with greater temporal precision. The 100 ms window was chosen based on our synaptic conductance measurements, which showed that preferred- and null-direction inhibition diverged clearly within this timescale across all plaid conditions (Fig. 2d). The 20 ms step approximates the 16.7 ms interval between presented movie frames, so that successive windows advance by about one frame and tile the stimulus. For each cell, we identified the movie segments that evoked reliable direction-selective responses by testing the response gDSI within each sliding window against a trial-shuffled null distribution. Windows with gDSI significantly above chance were classified as direction-selective windows for a given cell (permutation test, *P* < 0.05, 1,000 shuffles; Methods).

To determine which directions DSGCs compute during natural movies, we compared each cell’s preferred direction measured with drifting gratings to the VS/AM direction of the movies in each direction-selective window (Fig. 6a and 6c; Methods). Across cells and DS windows, the DSGC preferred directions remained clustered around the aligned VS/AM direction (Fig. 6b and 6d), with median population offsets of −8° for optic flow and −19° for object motion. Repeating the analysis with non-overlapping 100 ms windows yielded similar population offset distributions (−9° for optic flow and −18° for object motion), confirming that the results do not depend on window overlap.

**Fig. 6.**
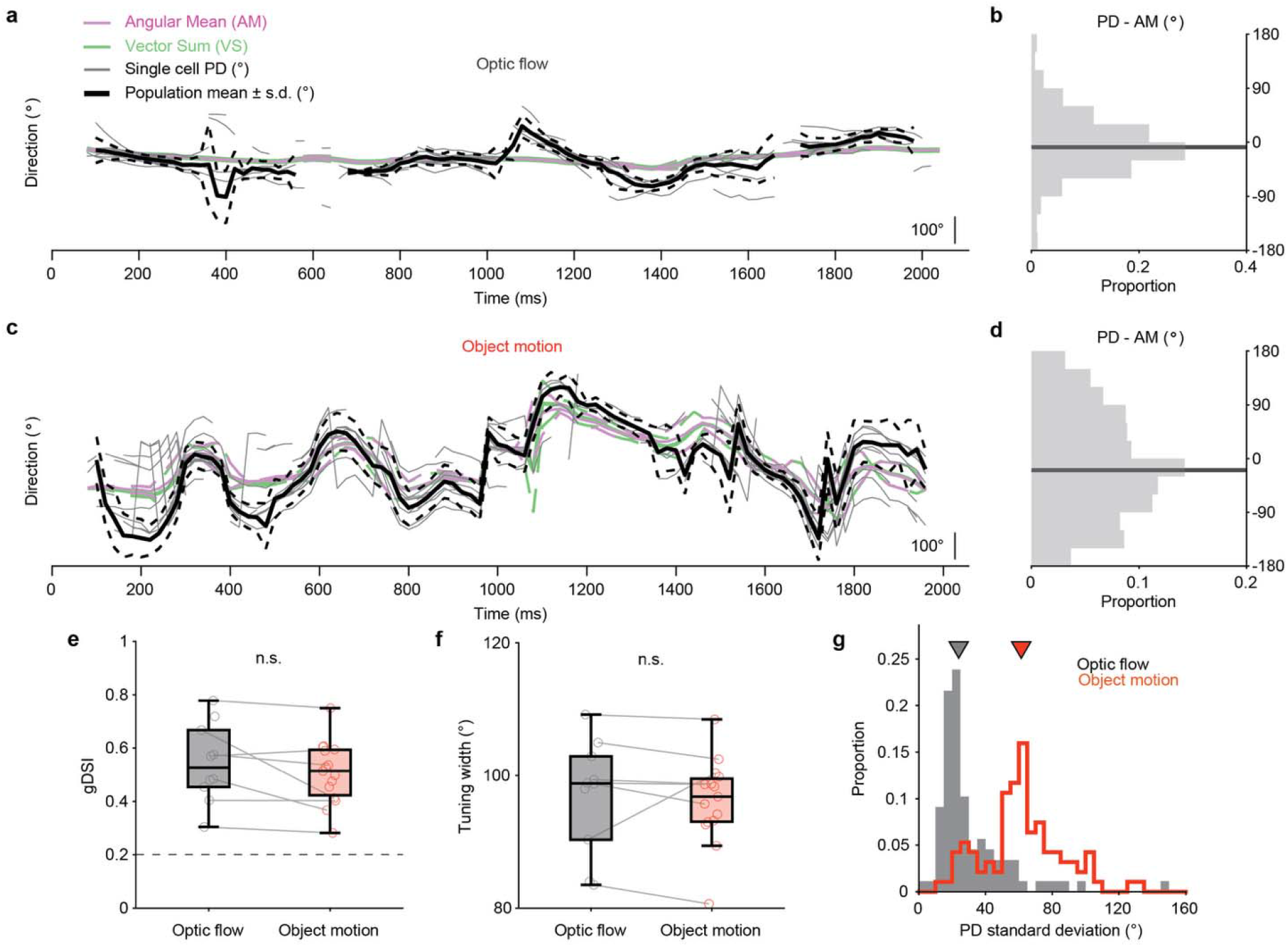
DSGCs encode the aligned VS/AM direction during natural movies. **a**, Time-resolved population preferred direction during optic flow, computed in sliding windows and plotted relative to each cell’s drifting-grating preferred direction. Gray traces show individual cells; solid and dashed black trace show population mean and mean ± s.d., respectively, when more than three cells contributed. **b**, Distribution of deviations between DSGC preferred direction and stimulus AM direction across all direction-selective windows during optic flow. **c**, Same as **a** for object motion. **d**, Same as **b** for object motion. **e**, Time-averaged tuning strength (gDSI) for each cell during optic flow and object motion. Median gDSI values were 0.53 for optic flow and 0.51 for object motion (*P* = 0.50, Student’s t-test). **f**, Time-averaged tuning width for individual cells (optic flow median 98° and object median 96°; *P* = 0.79, Student’s t-test). **g**, Distribution of preferred-direction standard deviation across direction-selective windows during optic flow (*n* = 88 windows) and object motion (*n* = 94 windows) movies. Inverted triangles indicate medians (24° for optic flow and 61° for object motion; *P* < 0.01). Sample sizes: *n* = 10 cells for optic flow and *n* = 15 cells for object motion throughout, of which 7 cells were tested with both movies (gray connecting lines in **e** and **f**).

### Wider angular range during object motion increases preferred direction variability

We then asked whether the broader angular range of object motion, relative to optic flow, affects DSGC tuning. Although object motion contains greater angular dispersion than optic flow (median angular range 52° for object motion vs 4° for optic flow), the frame-wise angular range still fell below 90° most of the time (Fig. 5b). DSGCs’ tuning strength and width were similar between the two stimulus classes (Fig. 6e and 6f). In contrast, object motion markedly increased the variability of preferred-direction angles across cells (Fig. 6g). This pattern is consistent with our plaid results: when angular range remained below 90°, DSGC tuning strength and width were largely preserved (Fig. 1e and f), whereas increased angular range resulted in greater variability in preferred direction (Fig. 1g). Thus, larger angular ranges can increase the variability of preferred directions at the population level without substantially altering the tuning strength of individual cells. Together, these results show that DSGCs remain tuned to the aligned VS/AM direction during natural motion, while broader angular range increases the variability of their preferred directions when sampled across the population.

## Discussion

Our results reveal how the canonical retinal direction-selective circuit computes motion direction for heterogeneous motion fields, a scenario visual animals often encounter in the natural environment. We show that the retina computes the AM of local motion vectors for both artificial and natural stimuli, and that the precision of direction signaling deteriorates with increasing angular dispersion of motion vectors (Fig. 7a). The computation of AM is implemented at the locus of inhibitory inputs from SACs to DSGCs. Importantly, natural motion stimuli differ from drifting plaids in a fundamental way: distinct vector computations, such as AM and VS, converge onto a consensus direction during natural movies. This convergence reduces ambiguity in local motion integration and simplifies direction estimation in the natural environment. Together, our study demonstrates the power of combining artificial and natural stimuli to uncover both the algorithm and ethological relevance of neural computation.

**Fig. 7.**
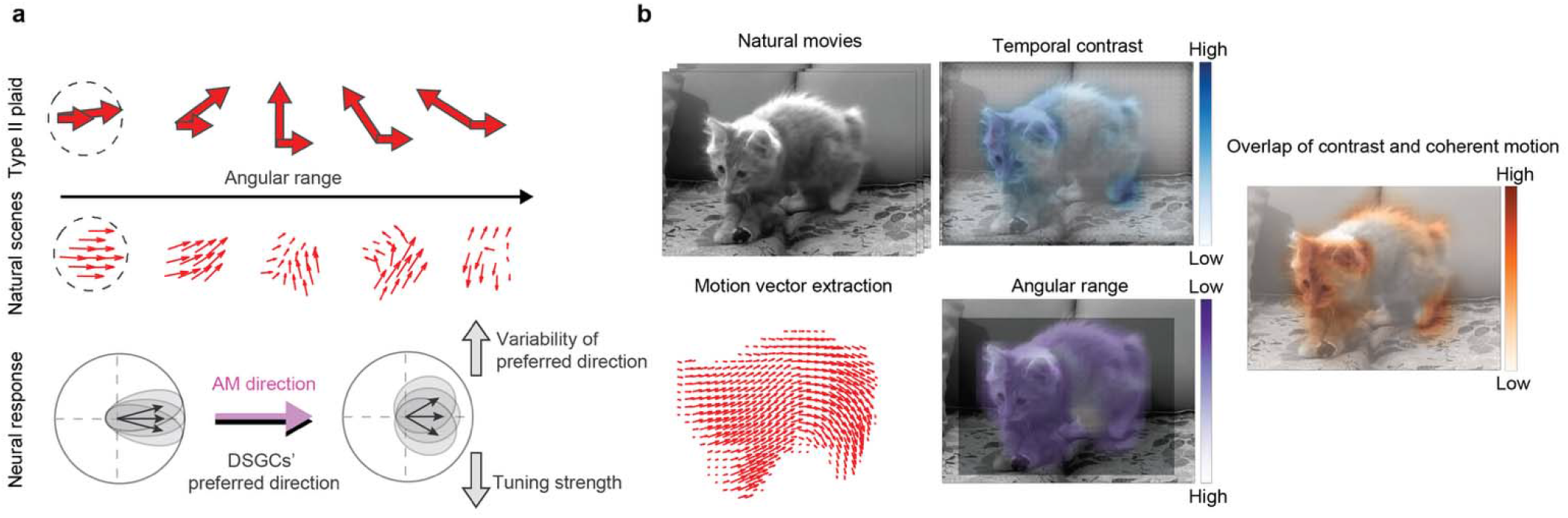
Summary of DSGC signaling during heterogeneous motion fields. **a**, As angular range increases for Type II plaids and natural stimuli, local motion vectors become more directionally heterogeneous. This causes more variable preferred directions across the DSGC population, and tuning strength decreases at obtuse angular ranges. **b**, DSGCs tune to the aligned AM/VS direction of natural movies. Preferred-direction firing rates depend on temporal contrast (temporal contrast color map), whereas tuning precision depends on motion coherence (angular range color map). Jointly, these tuning properties indicate that spatiotemporal regions of high temporal contrast and high motion coherence (right, combined color map) evoke strong, highly precise direction signaling in DSGCs (illustrated by Supplementary Video 3 and 4).

### Natural stimuli bypass algorithmic constraints in motion estimation

Our motion vector analysis reveals a remarkable simplicity beneath the apparent complexity of natural movies: distinct vector computations, such as angular mean and vector sum, converge onto a unified direction. This convergence stems from the spatiotemporal structure of the motion vector ensemble: natural motion vectors predominantly share similar speeds and angles, and occasional periods of large angular dispersion are transient and shorter than the motion integration window of the direction-selective circuit. This convergence persists across spatial scales larger than the receptive fields of DSGCs, suggesting that it is a broadly applicable principle for investigating direction selectivity in visual neurons beyond the retina, such as in the superior colliculus and visual cortex. Because of this convergence, natural stimuli eliminate the strict requirement for any specific vector computation to encode motion direction, thereby relaxing the evolutionary pressure to favor one algorithm over another. Instead, “all roads lead to Rome”: multiple computational algorithms can lead to the same solution.

The algorithmic convergence of natural motion vectors sheds new light on previous findings using artificial drifting plaids. Type II plaid stimuli provide a powerful means to probe the algorithm of motion integration because IOC, VS, and AM predict different directions for the same stimulus^32^. Studies using plaids have identified distinct algorithms of motion processing across different brain regions and species^50–54^. For example, while mouse On-Off DSGCs compute the AM direction, direction-selective cells in the mouse superior colliculus (SC) perform a probabilistically constrained VS computation: their preferred directions align with VS directions for plaids with 20°, 45°, and 160° cross angle, but shift toward IOC direction at 135° cross angle^53^. In tree shrews, which are more closely related to primates than to rodents, direction-selective cells in the SC compute the VS direction for plaids of all cross angles^54^. In the primate visual system, IOC is generally favored because it most closely matches human perception^30,32,52,55^. However, neural and perceptual responses consistent with all three vector computations have been reported^56–60^. Perceived motion direction depends on a number of factors such as relative contrasts^61^, spatial frequencies of the component gratings^30^, duration of stimulus presentation^57,59^, cross angle between the components^58^, eccentricity^57^, and aperture size^60^. The source of this variability in plaid response is not fully understood, but our study suggests that it might at least partially stem from the lack of algorithmic convergence of vector computations in the plaid stimulus. If different visual areas of an animal use different vector computations to integrate motion vectors, each area will compute a different direction for plaids, instead of a consensus direction for natural movies.

Distinct mathematical computations of motion vector integration that become less separable under natural vision may explain why multiple algorithms are used for computing direction selectivity in various brain regions and species. It may also shed light on why motion direction is typically perceived as clear and stable during natural vision despite the heterogeneity of local motion, while drifting sinusoidal plaids, which lack such algorithmic convergence, can generate unstable and ambiguous perception of direction. Future studies on motion processing in downstream visual areas and across species will further elucidate the implications of the convergence of natural motion vector computations beyond the retina.

### On-Off DSGCs are neither component-motion nor pattern-motion selective, but integrate motion vectors in a cross-angle-dependent manner

The directional tuning properties of On-Off DSGCs to Type II plaids indicate that these cells neither signal individual component motion vector directions nor the global motion direction of the plaid pattern. If DSGCs were tuned to component motion directions, we would expect the plaid tuning curve to be bimodal, with one peak when the slow component moves in the cell’s preferred direction and another peak when the fast component moves in the cell’s preferred direction, resulting in a monotonic decrease in direction selectivity and an increase in tuning width as cross angle widens, neither of which we observed (Fig. 1e and 1f). Although direction selectivity decreased at 160°, directional tuning curves remained unimodal (Extended Data Fig. 1f) and the population median gDSI remained above 0.2 (a conventional threshold for direction selectivity in the literature), indicating that DSGCs are tuned to an integrated motion direction even when the component motion directions are strongly divergent. Conversely, if the cells encoded the motion of the entire plaid texture, we would expect the preferred direction to align with IOC and no cross-angle dependent tuning: this is not what we observed either. Notably, 15° and 160° plaids have similar spatial structures (both are superpositions of two nearly parallel gratings) yet elicit markedly different preferred directions relative to their respective IOC directions in DSGCs. The two cross-angles also evoked DSGC responses of significantly different tuning strength and tuning width (Fig 1e and 1f), further indicating that DSGCs are not simply tuned to the motion direction of the overall plaid pattern. Together, these findings establish that On-Off DSGCs weight and integrate the two-vector stimulus in a cross-angle dependent manner, rather than encoding individual component motion directions independently or the overall direction of the plaid pattern. Therefore, motion vector integration begins at the earliest stage of visual processing in the retina.

### SACs perform vector integration

Directionally tuned inhibition from SACs is critical for retinal direction-selectivity^13,26^. Here we found that SAC-mediated inhibitory tuning also accounts for DSGC responses to plaids, suggesting that the vector integration is already implemented at the level of SAC outputs. Several lines of evidence support this idea. First, SACs have radially symmetric dendritic arbors that each prefers motion away from the soma^14,33,62–67^. Thus, plaids with small cross angles will activate largely overlapping branches, whereas plaids with large cross angles will recruit more distinct branches. More heterogeneous activation of the SAC dendritic tree might contribute to increased variability in IPSC preferred direction. Second, because the tuning and amplitude of inhibition are largely invariant to stimulus speed^21,33^, simultaneous activation of distinct SAC branches with similar synaptic weight could explain why inhibition to plaids is tuned to the angular mean direction. However, the deviation of preferred direction at obtuse cross angles remains unresolved, as do the loci of nonlinearity that keep tuning unimodal at large cross angles. Future studies examining how individual SAC dendrites are activated during heterogeneous motion will further elucidate the biophysical mechanisms of vector integration by SAC dendrites.

### On-Off DSGCs use the same vector integration rules to signal motion in Type II plaids and natural movies

Response properties of RGCs measured with carefully controlled artificial stimuli often differ from those measured with dynamic, complex natural stimuli. For example, the On-Off polarity of some RGCs can change depending on specific natural images or ambient luminance^68–70^, and the linearity of spatial integration in primate On parasol cells differs for sinusoidal gratings versus natural images^71,72^. These examples highlight the sophisticated contextual modulation of RGC responses that is highly sensitive to image statistics. In this study, we found an interesting counterexample: several aspects of On-Off DSGC responses are well preserved across artificial Type II plaids, optic flow, and object motion-dominated natural movies. For all three types of motion stimuli, 1) DSGCs remain unidirectionally tuned in heterogeneous motion fields; 2) Their preferred direction angles become more jittered when component motion vectors are more dispersed; 3) The temporal structure of preferred-direction firing is shaped by the temporal contrast of the stimulus. Therefore, despite pronounced differences in motion statistics, vector integration rules of DSGCs revealed by artificial stimuli are applicable to DSGC responses to natural movies.

### Single-neuron decoding constraints and future outlooks

Our finding that On-Off DSGC’s direction-selective spiking response is jointly modulated by motion vector coherence and temporal contrast highlights two constraints on decoding motion direction from single-neuron responses. First, in heterogeneous motion fields, the directional tuning of individual DSGCs loses precision and jitters around the angular mean (AM) direction, thereby degrading single-cell direction estimates. Second, because individual DSGC firing rates encode both direction and temporal contrast simultaneously, extracting one feature from the other introduces decoding ambiguities at the single-cell level.

Theoretical analysis indicates that pooling responses from the four subtypes of On-Off DSGCs that prefer four cardinal directions enables downstream decoders to extract precise motion direction, despite the broad tuning of individual cells and their sensitivity to other stimulus parameters^73^. Earlier work using moving bar stimuli in the rabbit retina demonstrates that while On-Off DSGC spike count varies with multiple stimulus parameters including direction, speed, and luminance, these parameters are multiplicatively separable through independent functions, suggesting a plausible population code to uniquely report direction^74^. The advantage of a population code is further demonstrated in the salamander retina, where ambiguities of luminance and motion information in individual DSGC responses to texture motion can be resolved by decoding population responses from cells preferring multiple directions^75^. Interestingly, a previous study indicates that luminance signals pose less of a challenge for direction encoding in mouse On-Off DSGCs, as their luminance responses are largely suppressed when natural movies span both the receptive field center and surround^76^. However, decoding motion direction from single-cell responses faces other constraints, such as response sparsity and preferred-direction variability, which can be mitigated by population decoding^77^. In this study, we restricted identical natural movies to the receptive field center of individual DSGCs and measured their responses using patch clamp recordings. Consequently, our data characterize single-cell response variability to a localized center stimulus, rather than capturing population-level features, such as signal correlations and trial-to-trial noise correlations, known to influence DSGC population decoding^40,77,78^. Future studies utilizing multielectrode array recordings or population calcium imaging, paired with natural movies spanning larger retinal fields, will be required to understand the coding performance of the On-Off DSGC ensemble for complex motion fields.

In summary, we used artificial plaid stimuli to reveal the computational logic of motion vector integration by the retina and used natural movies to reveal the demands of real-world motion detection. Understanding ethologically relevant computations of the retina is the first step in unraveling how the visual system solves the challenges of navigating the natural environment.

## Methods

### Animals

All procedures were approved by the University of Chicago Institutional Animal Care and Use Committee (protocol number ACUP 72247) and followed the NIH Guide for the Care and Use of Laboratory Animals and Public Health Service policy. Mice were housed on a 12 h light/dark cycle, in groups of two to five animals per cage, with food and water available ad libitum at room temperature and 40-60% humidity. Experiments used mice of both sexes at postnatal day 21-40. For two-photon calcium imaging, Vglut2-ires-Cre knock-in mice (JAX stock 028863; RRID: IMSR_JAX:028863) were crossed to the floxed GCaMP6f Ai95 reporter line (JAX stock 028865; RRID: IMSR_JAX:028865). For targeted electrophysiological recordings, we used the Drd4-eGFP mice, which was originally developed by MMRRC (http://www.mmrrc.org/strains/231/0231.html) in the Swiss Webster background and subsequently backcrossed to C57BL/6 background.

### Isolated retina preparation

Isolated retinas were prepared for both imaging and electrophysiology. Mice were dark-adapted for more than 30 min, anaesthetized with isoflurane and euthanized by decapitation. Under infrared illumination, retinas were isolated from the pigment epithelium in Ames’ medium (Sigma-Aldrich) bubbled with 95% O2/5% CO2 and cut into halves. Each retinal piece was mounted ganglion-cell-layer up on filter paper over an approximately 1.5 mm^2^ aperture, and cells near the center of the aperture were used for recordings or imaging. During experiments, retinas were continuously perfused with oxygenated Ames’ medium at 32-34°C and were kept in the dark except during visual stimulation and brief two-photon scans used for targeting and recording from fluorescent cells.

### Visual stimulation

Visual stimuli at low photopic range were generated on an OLED display (eMagin, 800 x 600 pixel resolution and 60 Hz refresh rate) via custom Python and MATLAB functions written with Psychophysics Toolbox^79^ and presented to the retina at a resolution of 1.1 or 1.5 µm/pixel focused onto the photoreceptor outer segments. Distances on the retina were converted to visual angle using 30 µm per degree. The positive-contrast moving bars were 220 µm (7°) wide and 1100 µm (37°) long and were used to identify On and Off response components; bars moved along their long axis in 12 pseudorandom directions at 440 µm/s (15°/s) with two repetitions per direction. Sinusoidal drifting gratings had a spatial frequency of 220 µm/cycle on the retina (∼ 0.14 cycles/°) and temporal frequencies of 1.5 and 4.5 Hz, corresponding to speeds of 330 and 990 µm/s (11 and 33 °/s). For each temporal frequency, gratings were presented for 2 s, followed by a 2 s gray screen, in 12 block shuffled directions separated by 30 degrees, with three repetitions per direction. Type II plaids were generated by superimposing two sinusoidal gratings moving at 11 and 33°/s at a defined cross-angle; for calcium imaging, cross-angles of 15°, 45°, 90°, 135° and 160° were used, and each composite plaid was presented for 2 s per trial, followed by a 2 s interstimulus interval, in 12 directions with three repetitions per direction. All stimuli moved within a 660µm-diameter aperture on the retina and cells near the center of the aperture were used for recordings to avoid edge effects.

For electrophysiology, all gratings and plaid stimuli were centered on the soma of the recorded cell and moved within a 300 µm diameter aperture. Slow and fast gratings had the same spatial and temporal frequency as those used for calcium imaging. Type II plaid cross-angles were 15°, 90° and 160°.

For spike recordings in response to natural movies, two movie stimuli were used: one dominated by optic flow and one dominated by object motion. We used the camera field of view to convert image pixels to degrees of visual angle. Movies were cropped to 10° in diameter on the retina, deliberately larger than the 7° receptive field center used for motion vector analysis, so that the stimulus covered the RF center of every recorded DSGC despite cell-to-cell variation in center size. Movies were converted to grayscale based on the color channels, rescaled to match the range of our OLED monitor, and presented on top of a uniform gray background. Each movie contained a 500 ms hold of the first movie frame, followed by the movie and a 500 ms hold of the last frame. The movie was repeated twice in 12 pseudo-randomly permuted rotations with 1 s of interstimulus gray screen in between. Movie frames were temporally resampled to the 60-Hz refresh rate of the OLED display using *numpy.interp()* so that image speeds on the retina matched those measured in the source clips. After interpolation, the optic-flow movie contained 115 movie frames and the object-motion movie contained 112 movie frames, corresponding to durations of 1.92 s and 1.87 s respectively.

### Two-photon calcium imaging of On-Off DSGCs

GCaMP6f fluorescence was imaged using a customized two-photon laser-scanning microscope (Bruker Nano Surfaces Division) described in^63^. Briefly, GCaMP6f was excited with a Ti:sapphire laser (Coherent, Chameleon Ultra II, Santa Clara, CA) tuned to 920 nm, and laser power was adjusted to avoid fluorescence saturation. Visual stimuli were presented after 10 s of continuous scanning to allow the laser-induced fluorescence transient to adapt to baseline. To separate stimulus light from GCaMP6f emission, the OLED output was passed through a blue band-pass filter (Semrock, Rochester, MA) peaking at 470 nm, and two custom notched filters (Bruker Nano Surfaces Division) were placed before the photomultiplier tubes to block light of the same wavelength. Time series of fluorescence were acquired through a 20x water-immersion objective at 10 Hz or faster. TTL pulses at the start and end of each stimulus direction were used to align imaging data to the visual stimulus. Responses to moving bars, gratings and plaids were acquired in separate time blocks, with at least 2 min between blocks to avoid photobleaching.

Analysis was performed using Suite2p^80^ and MATLAB R2024b. Soma positions were used to match the same cells across time blocks. Raw fluorescence signals were motion corrected with Suite2p, and somatic regions of interest (ROI) were manually curated in the Suite2p graphical interface. For each ROI, baseline fluorescence (F0) was the mean of the lowest 25% of raw fluorescence values (∼1 sec) during each 4-s trials. Each ROI was manually inspected and repetitions with unstable responses were excluded. (F-F0)/F0 traces were then clipped into individual trials, sorted by stimulus direction, and averaged over repetitions. Response magnitude was defined as the time integral of the mean ΔF/F0 trace across repetitions.

For each ROI, response reliability was quantified with a quality index^5^,

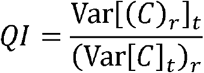

where *C* is the response matrix, *t* denotes time, and r denotes trial repetitions. ROIs with reliable grating responses (QI>0.12) were further analyzed to assess direction selectivity. We identified putative DSGCs as cells with gDSI ≥ 0.2 for either slow or fast gratings. To separate On–Off DSGCs from On-DSGCs, we manually inspected their moving bar responses and extracted cells that exhibited both ON and OFF responses to moving bars. These curated On-Off DSGCs were used for the subsequent Type II plaid calcium analyses.

### Tuning quantification

To quantify the strength of direction tuning, we calculated the global direction selectivity index (gDSI) as the magnitude of the normalized complex sum from responses to stimuli in each of the 12 stimulus directions evenly spanning 360°:

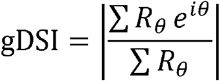

Responses *R_θ_* at stimulus direction *θ* (in radians) were measured as the integrated ΔF/F0 for calcium recordings, the integrated conductance for voltage clamp recordings, and the total spike count for spike recordings. The preferred direction was the angle of the resultant vector in degrees:

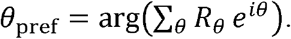

Tuning width was estimated by fitting a von Mises distribution ^81^ and calculating the full width at half height (FWHH) of the fitted curve.

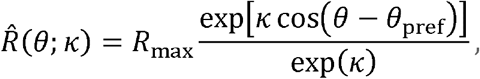

Where *R_max_* is the maximum response, *θ_pref_* is the preferred direction in radians, and *K* is the concentration parameter for the width of the direction tuning.

### Calculating the preferred direction for Type II plaids and aligning across cells

Plaids with different cross-angles were analyzed separately. For each Type II plaid, we compared three candidate motion-direction estimates: angular mean (AM), the circular average of the two component directions independent of speed; vector sum (VS), the speed-weighted mathematical sum of the component velocity vectors; and intersection of constraints (IOC), the predicted plaid pattern-motion direction defined by the intersection of constraint lines perpendicular to each component vector.

For each cell, we first calculated the preferred plaid stimulus from the plaid tuning curve. For a given plaid cross-angle with fixed speed ratio (1:3), each pair of component directions uniquely defines the corresponding AM, VS, and IOC directions, with fixed angular offsets among the AM, VS and IOC directions.

To determine which direction a cell prefers, we computed the circular difference between each cell’s grating preferred direction (PD) and the AM, VS and IOC directions of its preferred plaid stimuli. For each plaid cross-angle, plaid gDSI, tuning width and preferred-direction variability were then summarized across cells. Population variability in preferred plaid direction was quantified as the circular standard deviation.

### Cell-targeted patch-clamp electrophysiology

Targeted electrophysiological recordings were made from GFP-labelled On-Off DSGCs in Drd4-GFP retinas according to ^82^. GFP-positive cells were targeted with brief two-photon imaging at 920 nm, and DSGC identity was confirmed physiologically from responses to drifting gratings. Recordings were acquired with pCLAMP 10 software, a Digidata 1550A digitizer and a MultiClamp 700B amplifier (Molecular Devices), low-pass filtered at 4 kHz and digitized at 10 kHz.

For whole-cell voltage-clamp recordings, patch electrodes of 3-5 MΩ were filled with a cesium-based internal solution containing 110 mM CsMeSO4, 2.8 mM NaCl, 4 mM EGTA, 5 mM TEA-Cl, 4 mM Mg-ATP, 0.3 mM Na-GTP, 20 mM HEPES, 10 mM phosphocreatine, 5 mM QX314 and 0.025 mM Alexa 594, adjusted to pH 7.25. EPSCs and IPSCs were isolated by holding cells at command potentials corresponding to approximately -65 mV and 0 mV, respectively, after correcting for an approximately 10 mV liquid junction potential. For cell-attached spike recordings, electrodes of 3.5-5 MΩ were filled with Ames’ medium.

### Synaptic conductance analysis

Synaptic currents recorded in voltage-clamp recordings were analyzed using custom MATLAB scripts. For each sweep, the stimulus-evoked synaptic current was calculated by subtracting the mean pre-stimulus current from the raw trace. Baseline-subtracted currents were converted to conductance using reversal potentials of 0 mV for excitation and -65 mV for inhibition. For each cell, conductance traces were averaged across repetitions for each stimulus direction, and response magnitude was defined as the integrated conductance during the motion-evoked response window. To focus on the steady-state motion response, the first 200 ms after stimulus onset was excluded, and conductance was integrated over the remaining 1.8 s stimulus period.

For gratings and plaid responses, preferred direction and gDSI were calculated from the integrated conductance across directions as described above. IPSC grating preferred direction was defined as the slow-grating IPSC preferred direction. For the same cell, EPSC preferred direction was defined as 180° away from the slow-grating IPSC preferred direction. For Type II plaids, conductance responses were analyzed separately for each cross-angle as specified above. Preferred and null directions were taken from the plaid-evoked IPSC tuning measured in the same cell at the same cross-angle.

### Temporal contrast analysis

Temporal contrast was computed from stimulus frame-to-frame luminance changes. Within a given DSGC receptive-field center, the luminance changes within each subunit *b* were calculated as:

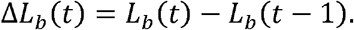

Positive and negative luminance changes were half-wave rectified into ON and OFF components and smoothed with a boxcar window,

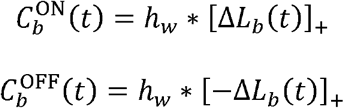

where [*x*]^+^ = max(*x*, 0), *h_w_* is a 100 ms boxcar window, and * denotes convolution. contrast available to the DSGC was then calculated by averaging the summed ON and OFF signals The total temporal across subunits (each 2° in diameter with 1° offsets, *N_b_* denotes the number of local subunits):

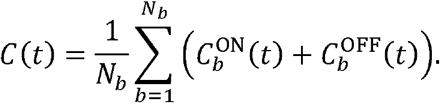

Temporal contrast was computed within a circular aperture matched to the DSGC receptive field center. For Type II plaid stimuli, we used a 7° diameter aperture. To compare temporal contrast with synaptic currents, the temporal contrast trace was shifted by 40 ms to account for synaptic latency, which was estimated from recorded conductance responses. For natural movies, temporal contrast was computed using the same approach within a 10° diameter aperture, matched to the size of the cropped movies. Temporal contrast traces were normalized within each movie and shifted by 100 ms to account for the spike latency estimated from the recorded spike responses to natural movies.

### Natural movie selection

Movie clips were selected based on their ethological relevance alone to span the types of visual motion most relevant to the mouse observer. Most of the object-motion movies were filmed for this study by the authors and an acknowledged contributor; the remainder were obtained from publicly available online recordings. The optic-flow movies were taken from a published open-source database^38^. Selection was completed before any motion vector extraction was performed, and no clips were excluded after analysis.

### Motion vector extraction from natural movies

Motion vector extraction was performed using custom Python and MATLAB scripts. Pixel-wise motion fields were estimated between consecutive movie frames using RAFT (Recurrent All-Pairs Field Transforms)^42^, a deep recurrent neural network for dense frame-to-frame displacement estimation. Briefly, RAFT first extracts learned visual features from each frame, then constructs multi-scale all-pairs correlation volumes that encode the similarity between pixels across the two frames. A recurrent operator then iteratively refines a dense displacement field, allowing RAFT to capture both large image shifts and fine local motion. We used the pretrained *raft-things* model to estimate motion in optic-flow and object-motion movies. The resulting motion fields were visually inspected and checked for accuracy in direction and speed by manually calculating direction and speed of sampled moving objects in the movie and comparing them with RAFT measurements. Comparison with the Farneback method using *cv2.calcOpticalFlowFarneback()* confirmed that RAFT more reliably separated moving objects from background motion in these natural movies.

Pixel-wise motion vectors were first converted from pixels per frame to degrees per second using the movie frame rate and the visual angle of the camera’s field of view. Motion vectors were then filtered to speeds between 5 and 60 °/s, corresponding to the speed range over which DSGCs showed robust direction-selective responses^21^. To approximate spatial pooling of bipolar inputs to the ganglion cell, retained vectors were averaged within circular bipolar-cell-sized apertures, yielding one component motion vector per bipolar cell unit. These local motion vectors were then pooled within the DSGC receptive field. For each DSGC receptive field across consecutive frames, we computed five summary statistics: the vector-sum (VS) direction from the speed-weighted component vectors, the angular-mean (AM) direction from the circular mean of local directions, the mean speed across local vectors, the speed ratio as the ratio of the maximum to minimum local vector speed, and the angular range as the smallest circular arc containing all local directions. The same analysis was repeated using non-overlapping apertures of 7°, 15°, and 30° diameters to examine how motion statistics changed with receptive-field size.

### Dynamics of preferred-direction firing rate during natural movies

To estimate the dynamics of preferred-direction firing during natural movie stimulation, spikes from each cell were pooled across repetitions for each movie rotation and binned at 1 ms resolution. Trial-averaged firing rates for each movie rotation were estimated by smoothing the binned spike trains with a Gaussian kernel with σ = 30 ms. The preferred-direction firing-rate envelope was defined as the maximum smoothed firing rate across the 12 movie rotations at each time point. For comparison across cells, each cell’s envelope was normalized to its peak firing rate. Population traces were then computed as the mean normalized preferred-direction envelope across cells.

### Calculating the preferred direction for natural movies and aligning across cells

To estimate which directions DSGCs prefer during natural movies, we first measured each cell’s preferred direction from responses to drifting gratings,θ*_i_* for cell *i.* Spike responses during natural movies were then analyzed using 100 ms sliding windows advanced in 20 ms steps. Within each window, spike counts were averaged across repetitions for each of the 12 movie rotations.

To identify windows in which a cell was significantly direction selective, we performed a permutation test ^5^. Spike counts were randomly shuffled across movie rotations and repetitions 1,000 times to generate a null distribution of gDSI. Windows were classified as direction-selective when the observed gDSI exceeded the shuffled null distribution at *P* < 0.05. Non-significant windows were excluded from subsequent analyses.

For each direction-selective window, we first computed the preferred movie rotation, *x_i_(t)* and then compared the stimulus VS and AM directions, *θ_vs_(ts)* and *θ_AM_(ts)* with the cell’s grating preferred direction, *θ_i_*. An offset near zero indicates that the cell fired most strongly to the movie rotation in which the VS or AM direction was aligned with its grating preferred direction.

### Statistical analysis

Statistical analyses were performed in MATLAB. Statistical methods were selected according to the experimental design and are reported in the corresponding figure legends. Unless otherwise indicated, individual data points represent cells, frames, or time bins as specified in each figure. Unless otherwise stated, data are summarized as mean ± standard deviation or median with interquartile range, as indicated in each figure. Circular variables were analyzed using circular statistics. Statistical significance was defined as *P* < 0.05. Nonsignificant comparisons are labeled n.s.; *, **, and *** indicate *P* < 0.05, *P* < 0.01, and *P* < 0.001, respectively. Exact tests, sample sizes, and *P* values are provided in each figure legend.

## Author contributions

W.W. and Z.D. designed experiments, filmed object-motion footage, and wrote the manuscript. Z.D. performed experiments and data analysis.

## Data availability

The data analyzed in this study will be posted to FigShare (www.figshare.com) at the time of publication.

## Code availability

Code used for analysis will be made available in a public GitHub repository upon publication.

## Acknowledgements

We thank Chen Zhang for managing the mouse colony; Stephanie E. Palmer and John H.R. Maunsell for helpful discussions; Matthew Ahmon for help with preprocessing the calcium-imaging data presented in Fig. 1; Changyu Sun for filming the cat-grooming clips; and members of the Wei Lab for careful reading of the manuscript and helpful feedback. This work was supported by NEI R01EY035268 and the McKnight Scholarship Award to W.W., and the Pritzker Fellowship and NIDA Training Grant R90DA 060338 to Z.D.

---

**Extended Data Fig. 1.**
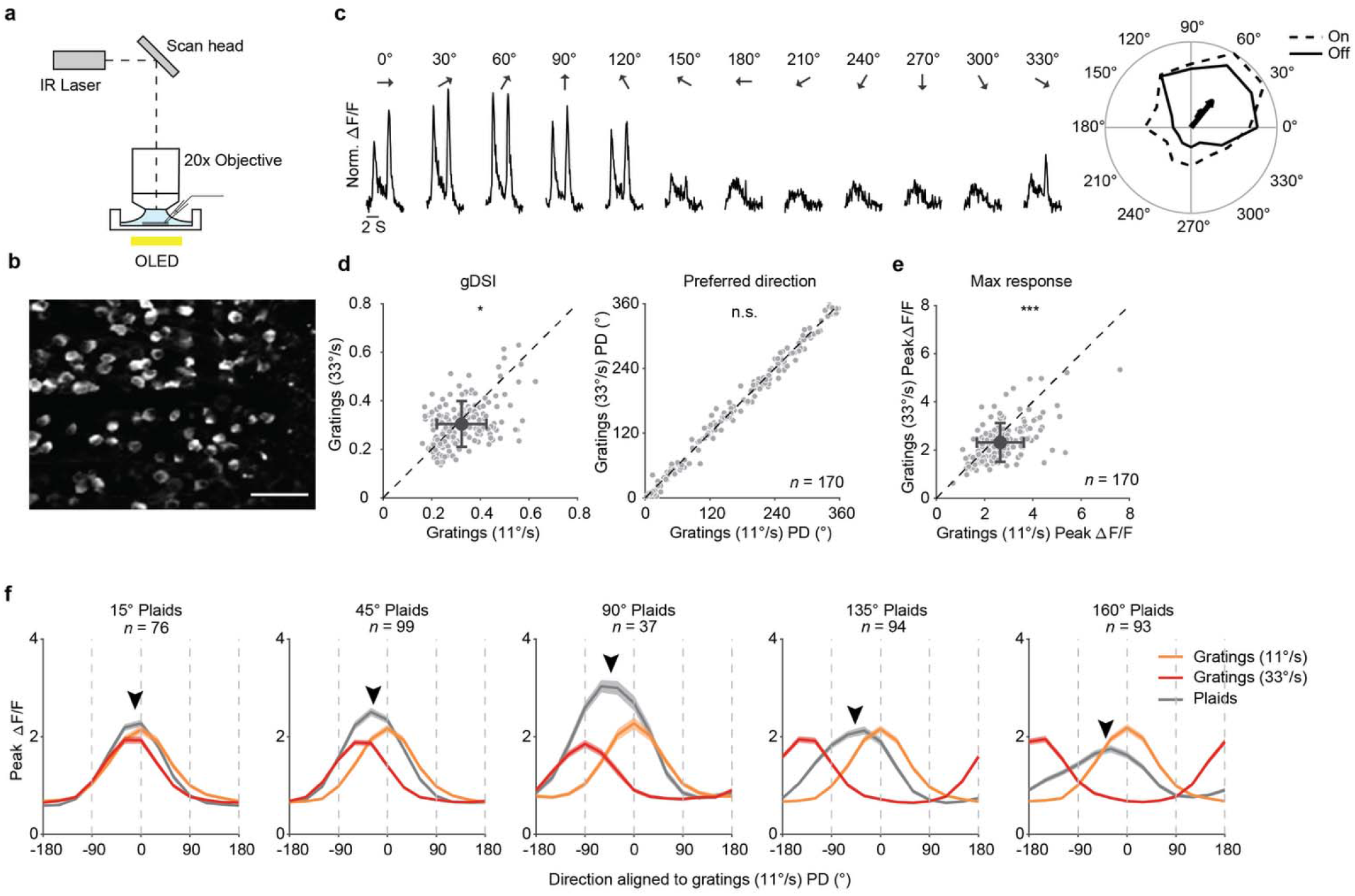
On-Off DSGCs exhibit unimodal direction tuning to gratings and Type II plaids. **a**, Schematic of two-photon calcium imaging. Visual stimuli are presented on an OLED display and focused on the photoreceptor outer segments of an isolated whole-mount retina. **b**, Representative standard-deviation projection of GCaMP6f fluorescence in the ganglion cell layer. Scale bar, 50 μm. **c**, Left, calcium responses of an example On–Off DSGC (same cell shown in Fig. 1c) to moving bars in different directions. Columns indicate stimulus direction. Right, moving bar tuning curves for On and Off responses. **d**, Left, comparison of tuning strength, measured as gDSI, for slow (11°/s) versus fast (33°/s) gratings. *n* = 170 cells, *P* = 0.03, Wilcoxon signed-rank test. Right, comparison of preferred directions for the same cells; *P* = 0.14, Wilcoxon signed-rank test. **e**, Comparison of peak response amplitude to slow (11°/s) versus fast (33°/s) gratings; *n* = 170 cells; *P* < 0.001, Wilcoxon signed-rank test. **f**, Population tuning to Type II plaids with cross angles of 15°, 45°, 90°, 135° and 160°. Responses were aligned to each cell’s slow grating (11°/s) preferred direction, with positive angles denoting counterclockwise shifts. Orange and red curves show responses to 11°/s and 33°/s gratings respectively; black curves show plaid responses. Arrowheads indicate population preferred directions: −11°, −26°, −42°, −47° and −44° for 15°, 45°, 90°, 135° and 160° plaids respectively.

**Extended Data Fig. 2.**
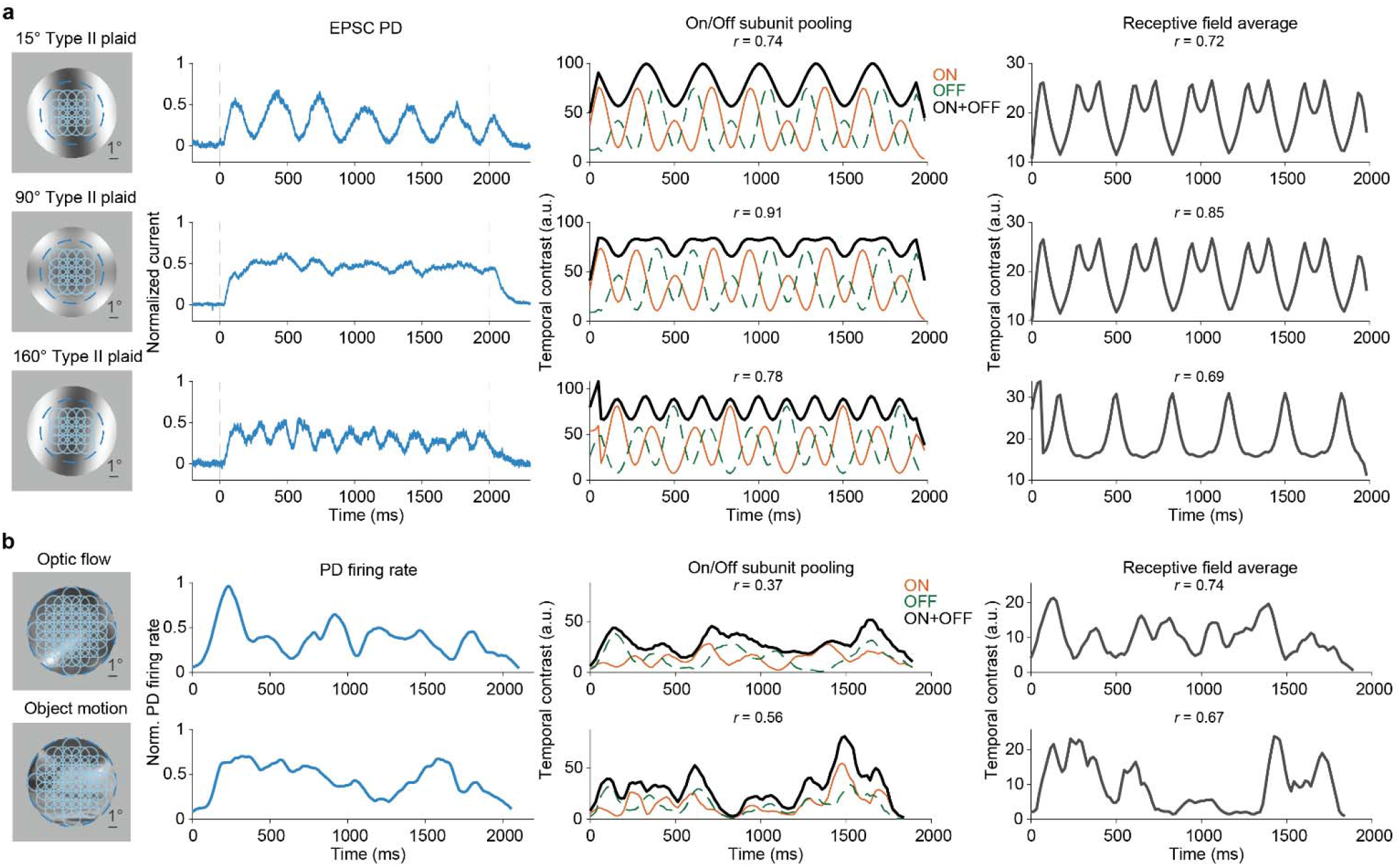
Temporal contrast calculated from ON/OFF bipolar subunit pooling differs from that calculated from receptive field averaging for Type II plaid and natural movies. **a**, Comparison of normalized EPSCs with the temporal contrast of Type II plaid stimuli at three plaid angles: 15°, 90°, and 160°. From left to right: example stimulus frames with bipolar subunits overlaid; normalized mean EPSCs in the preferred direction (PD) from data; temporal contrast of the stimulus computed from subunit pooling, with ON subunit activation shown in orange and OFF subunit activation in dashed green, and ON+OFF in black; temporal contrast computed from averaging luminance changes across the DSGC receptive field (7° diameter). Pearson *r* between normalized temporal contrast and EPSC PD is shown. **b**, Comparison of normalized preferred-direction firing rate with the temporal contrast of optic flow and object motion movies computed using the same ON/OFF subunit pooling and receptive field averaging methods as in **a**. From left to right: example movie frames with bipolar subunits overlaid; normalized preferred-direction firing rate; temporal contrast computed from local ON/OFF subunit pooling; temporal contrast computed from averaging luminance changes across the entire DSGC receptive field (10° diameter). Pearson *r* between normalized temporal contrast and PD firing rate is shown.

**Extended Data Fig. 3.**
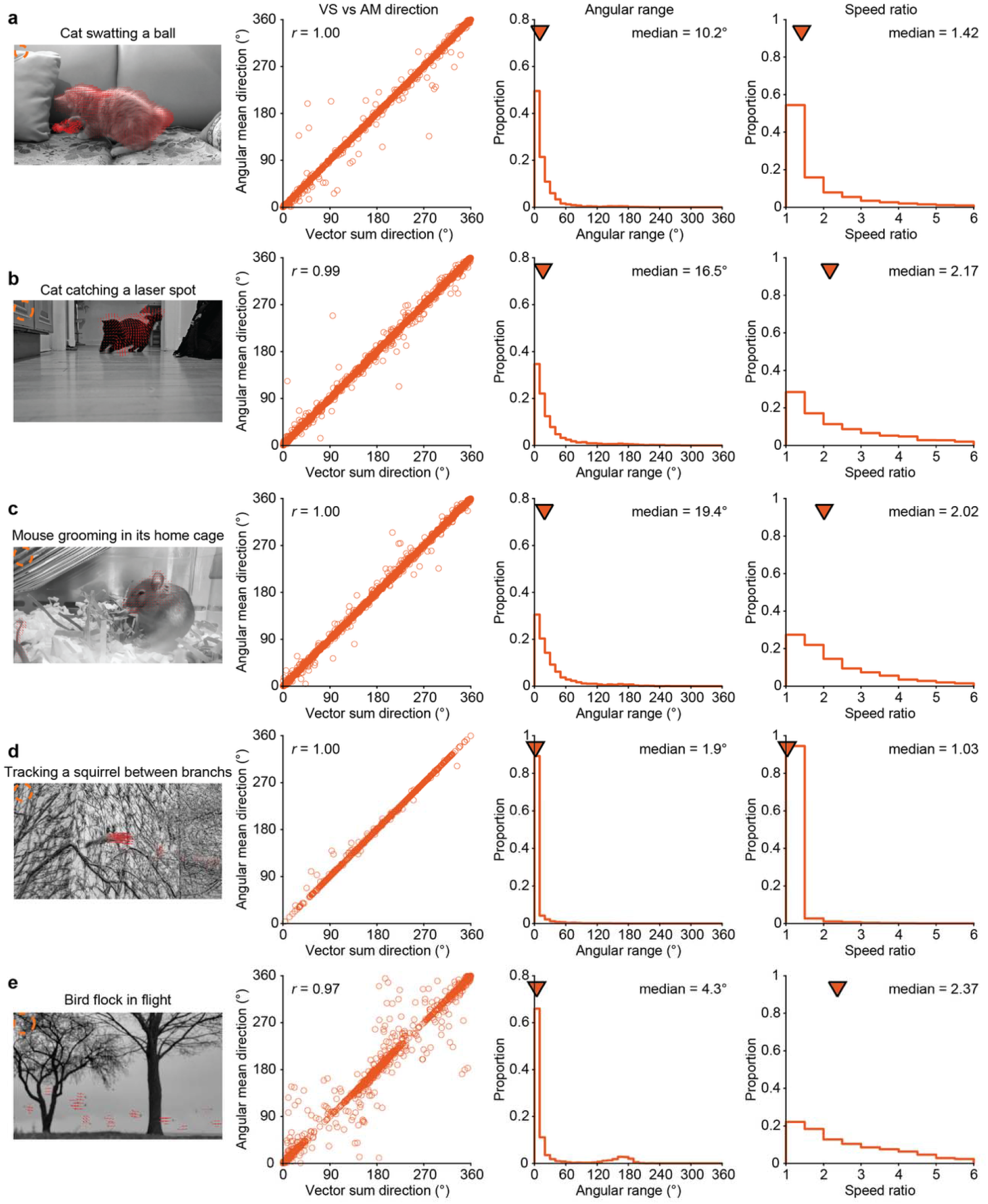
Motion statistics of example object motion movies. **a**, Left, representative frame from a home cat video. The orange dashed circle indicates the DSGC RF (7° diameter), and the red arrows indicate local motion vectors within bipolar RF units (2° diameter with 1° offsets). Center-left, comparison of VS and AM directions across all DSGC RFs and consecutive movie frames. Each point represents one RF across consecutive frames. *r* denotes Pearson correlation coefficient. Center-right, distribution of angular ranges for the same RFs across the movie clip. Rightmost, distribution of speed ratios across all RFs. Inverted triangles indicate the medians. **b**-**e**, same analyses as in **a** but for different movie clips.

**Extended Data Fig. 4.**
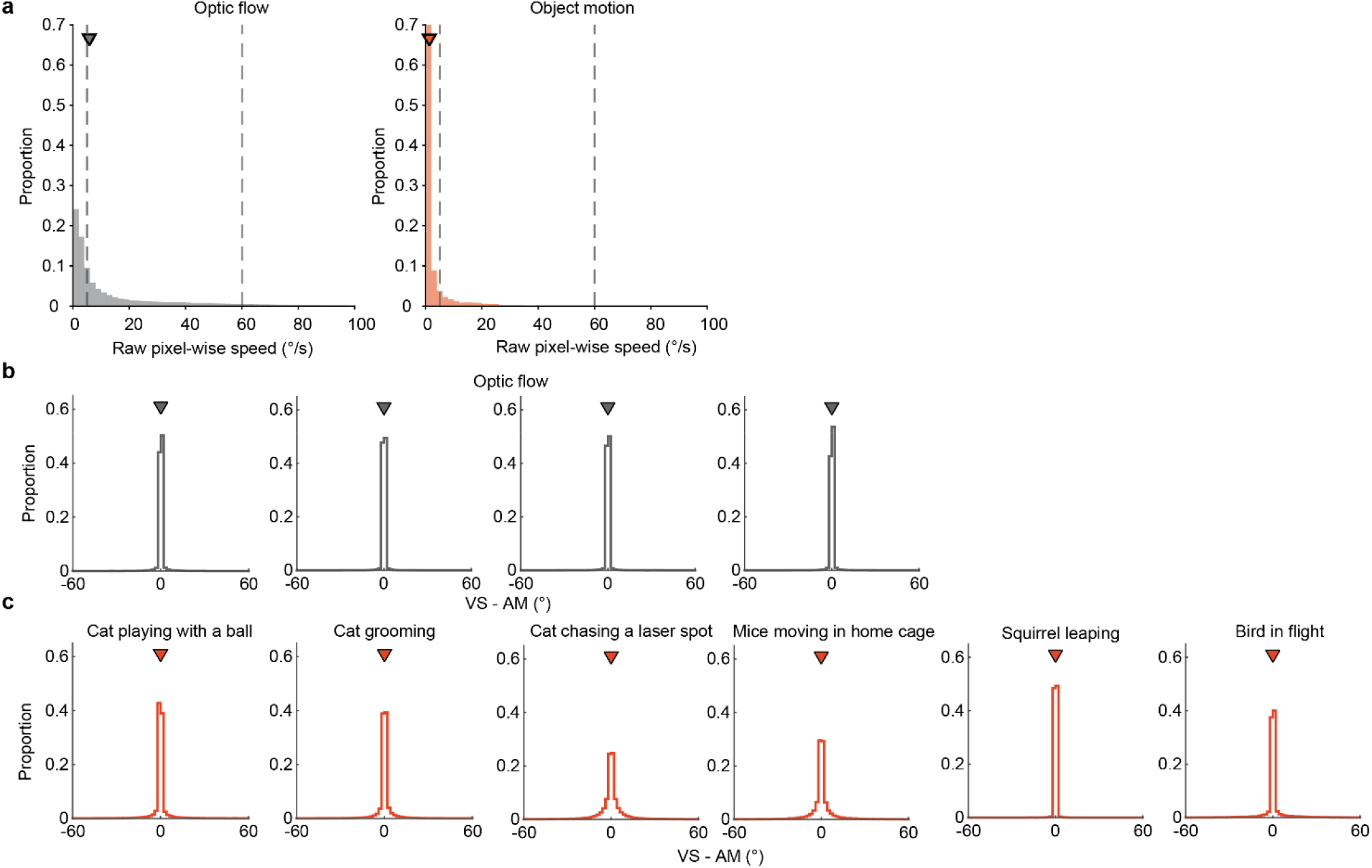
Raw pixel-wise motion statistics for optic-flow and object-motion movies. **a**, Distributions of raw, unfiltered pixel-wise motion speed for optic-flow movies (gray; *n* = 4 clips) and object-motion movies (orange; *n* = 16 clips). Inverted triangles indicate medians; dashed vertical lines mark 5 and 60°/s, the speed thresholds used in the filtered analyses. **b**, Distribution of VS-AM offsets computed from raw pixel-flow vectors from the same optic-flow movies analyzed in **a** with no speed filtering. **c**, Distribution of VS-AM offsets computed from the same object motion movies analyzed in **a** using raw pixel-flow vectors with no speed filtering. Object motion movies were pooled by behavioral motifs: cat playing with a ball (*n* = 2 clips), cat grooming (*n* = 4 clips), cat chasing a laser spot (*n* = 3 clips), mice moving in home cage (*n* = 4 clips), squirrel leaping (*n* = 2 clips), and bird in flight (*n* = 1 clip). For **b** and **c**, inverted triangles mark medians. All VS-AM medians were 0°.

## Supplementary videos

**Supplementary Video 1 | Optic-flow stimulus with local motion vectors.**

The optic-flow movie as presented to the retina. Red arrows indicate local component motion vectors within the cropped region. Playback is at 30 frames per second, half the 60 Hz stimulus presentation rate (0.5 × real time).

**Supplementary Video 2 | Object-motion stimulus with local motion vectors.**

As in Supplementary Video 1, for the object-motion movie as presented to the retina. Playback is at 30 frames per second, half the 60 Hz stimulus presentation rate (0.5 × real time).

**Supplementary Video 3 | Example object-motion movie.**

An uncropped object-motion movie shown as a reference for Supplementary Video 4. Playback is 10 frames per second; the original recording was 25 frames per second (0.4 × real time).

**Supplementary Video 4 | High temporal contrast and motion coherence regions in example object-motion movie.**

The same movie segment as in Supplementary Video 3, with regions of high temporal contrast and high motion coherence overlaid in orange.

